# Mechanisms of low susceptibility to the disinfectant benzalkonium chloride in a multidrug-resistant environmental isolate of *Aeromonas hydrophila*

**DOI:** 10.1101/2023.03.10.532024

**Authors:** Luz Chacón, Benno Kuropka, Enrique González-Tortuero, Frank Schreiber, Keilor Rojas-Jiménez, Alexandro Rodríguez-Rojas

## Abstract

Excessive discharge of quaternary ammonium disinfectants such as benzalkonium chloride (BAC) into aquatic systems can trigger several physiological responses in environmental microorganisms. In this study, we isolated a low susceptible strain of *Aeromonas hydrophila*, namely INISA09, to from a wastewater treatment plant in Costa Rica. We characterized its phenotypic response upon exposure to three different concentrations of BAC and determined the primary mechanisms related to its resistance using genomic and proteomic tools. The genome of the strain is 4.6-Mb with 4,273 genes, and we found a propound genome rearrangement and thousands of missense mutations when compared to the reference strain *A. hydrophila* ATCC 7966. We identified 15,762 missense mutations mainly associated with transport, antimicrobial resistance, and outer membrane proteins. In addition, quantitative proteomic analysis revealed the significant upregulation of several efflux pumps and the downregulation of porins when the strain was exposed to three BAC concentrations. Other genes related to membrane fatty acid metabolism and redox metabolic reactions also showed an altered expression. Our findings suggest that the response of *A. hydrophila* INISA09 to BAC primarily occurs at the membrane level, which is BAC’s primary target. This study sheds light on the mechanisms of antimicrobial susceptibility in tropical aquatic environments against a widely-used disinfectant, using an environmental strain that could serve as a new study model.

## 1 Introduction

Antimicrobial agents used for disinfection in domestic, industrial, and medical settings can enter wastewater and be released into the environment after treatment (Tezel and Pavlostathis, 2015). One of the most used disinfectants is the quaternary ammonium compound benzalkonium chloride (BAC), a cationic surfactant with biocidal properties against viruses, bacteria, and some fungi, including yeast. BAC has homologs with varying alkyl chain lengths and is used in large quantities, with over 450,000 kg manufactured or imported to the United States annually. As a result, the Environmental Protection Agency (EPA) has placed BAC on its High Production Volume list (Barber and Hartmann, 2022). During the COVID-19 pandemic, the consumption of quaternary ammonium compounds (QAC) products containing BAC increased significantly due to their widespread use. Over one million pounds of BAC are manufactured or imported to the United States annually. Therefore, the EPA has included BAC on its High Production Volume list (Barber and Hartmann, 2022). Remarkably, the mass load of QAC compounds in wastewater increased by 331% compared to pre-pandemic levels (Mohapatra et al., 2023).

BAC can remain in biosolids, such as activated sludge, for extended periods because a significant proportion of it gets integrated into the biomass, with around 95% of the compound adsorbed in just a few hours (Zhang et al., 2011). Furthermore, BAC degradation in solids is approximately 20 times slower than in the liquid phase, exacerbating the situation (Zhang et al., 2011; Barber and Hartmann, 2022). Previous studies have demonstrated that BAC is present in digested and sewage sludges used for fertilization purposes as land biosolids (Östman et al., 2017; U.S. Environmental Protection Agency, 2018). The biocidal effect of BAC begins when it interacts with cell membranes, leading to perturbation of the lipid bilayer and a progressive leakage of cytoplasmic materials outside the cell. BAC can also bind to anionic sites on surface membranes and induce osmoregulatory losses. BAC can affect bacteria’s respiration process, solute transport, and cell-wall biosynthesis at high concentrations. One common mechanism of microbial death is the solubilization of the cell membrane, which leads to the release of cellular contents into the environment (Gilbert and Moore, 2005). Although QACs are commonly associated with damaging cell membranes, recent research suggests their effects may extend beyond the surface to intracellular targets (Knauf et al., 2018). To survive under BAC exposure, some intrinsic responses have been described in Gram-negative bacteria, including structural cell membrane modification, efflux pumps activity to extrude biocides, biofilm formation, and formation of persister cells, and some of these adaptations can be related to cross-tolerance to other types of QAC and antibiotics (Mohapatra et al., 2023).

The most common bacterial models used to study the development of resistance to BAC are reference strains of *Pseudomonas aeruginosa* (Loughlin et al., 2002; Edirsana et al., 2014; El-Banna et al., 2019), isolates from food of *Listeria monocytogenes* (Romanova et al., 2006; Conficoni et al., 2016; Yu et al., 2018), and other clinically relevant isolates (Mahzounieh et al., 2014; Worthing et al., 2018; Abdelaziz et al., 2019). However, it is recognized that antimicrobial resistance originates in the environment, where waters, soil, and other sites can provide an unmatched gene pool with a great diversity that can exceed that of the animal and human microbiota. Consequently, antimicrobial pollution contributes to the mutation-based evolution of resistance (Larsson and Flach, 2022). For this reason, it is necessary to address the resistance profile and response to antibiotics and biocides to understand their impact on relevant bacterial species. Hypothesis-independent OMICs approaches allow the inference of the molecular mechanisms underlying antimicrobial resistance in such environmental bacteria without building these strains into elaborate model systems.

During a microbiological sampling campaign at a wastewater treatment plant near San José, Costa Rica, we identified a bacterial environmental isolate as *Aeromonas hydrophila* through sequencing. We designated this strain as *A. hydrophila* INISA09. *A. hydrophila* belongs to the Phylum Proteobacteria, Class Gammaproteobacteria, Order Aeromonadales, and Family Aeromonadaceae. It is a Gram-negative bacillus that is oxidase and catalase-positive, a glucose fermenter, and can reduce nitrates to nitrites. This bacterium is ubiquitous and typically present in aquatic environments (Fernández-Bravo and Figueras, 2020). *A. hydrophila*’s ability to colonize different natural and urban aquatic ecosystems makes it a desirable candidate for evaluating the dissemination of antimicrobial resistance (Igbinosa and Okoh, 2012; Karkman et al., 2018).

This study aims to characterize a naturally occurring bacterial isolate, a strain of *A. hydrophila* named INISA09, from a tropical activated sludge and understand its response to increasing benzalkonium chloride (BAC) concentrations using a quantitative label-free proteomics approach.

## 2 Materials and Methods

### 2.1 Bacteria and growth conditions

An *Aeromonas hydrophila* INISA09 strain, isolated an activated sludge in Costa Rica, was used as a bacteria model for all experiments with BAC. The isolate was recovered after an activated sludge exposition to BAC (Chacón et al., 2021) in trypticase soy agar (TSA) (Oxoid®, United States) enriched to 25 mg/L BAC (≥95.0% Fluka 12060 Sigma, United States). The *16S rRNA* gene was sequenced to identify the bacterial species. LB Lennox medium (Carl Roth, Germany) was used for all experiments. The growing temperature in all experiments was 30 °C, and when shaking conditions were needed, the shaker incubator was set to 200 rpm.

### 2.2 Genome sequencing

DNA extraction was conducted with DNeasy Blood & Tissue Kit (Qiagen, USA) using the pre-treatment step for Gram-negative bacteria recommended by the manufacturer. Illumina Hiseq platform in Novogene (Hong Kong) was used for DNA sequencing. Sequence quality was examined with FastQC (Brown et al., 2017). Low-quality reads and adapter removal from the paired ends for all sequences were filtered by Trim Galore v. 0.6.6 (Krueger, 2020) using a Phred quality cutoff of 30. Then, human reads were discarded by mapping them against the Genome Reference Consortium Human Build 38 (GRCh38; (Schneider et al., 2017)) using Bowtie2 v. 2.4.2 (Langmead and Salzberg, 2012) with default parameters. The reads were mapped against the different sequenced *A. hydrophila* strains (Table S2) using Bowtie2 with default parameters to remove all potential bacterial contamination. After that, for the de novo assembly, Shovill v. 1.1.0 (Seemann, 2020) was executed using SPAdes v. 3.15.0 (Bankevich et al., 2012) as the main assembler and with default parameters. The resulting contigs were scaffolded against *Aeromonas hydrophyla* ATCC 7966 using CSAR (Chen et al., 2018). Duplicated entries were removed using SeqKit v. 2.3.1 (Shen et al., 2016). All the resulting scaffolds were annotated with Prokka v.1.14.6 (Seemann, 2014) using the bacterial genetic code for the gene and tRNA prediction (-gcode 11), allowing the search of ncRNA elements (-rfam), predicting the Gram-negative signal peptides in the CDSs (-gram neg), and complaining with the GenBank specifications for the file formatting (-compliant), a Average Nucleotide Identity (ANI) (Jain et al., 2018) and Snippy v. 4.6.0 (Seemann, 2015) was used to detect the conservation degree and genomic variation, respectively, between *A. hydrophila* INISA09 and *A. hydrophila* ATCC 7966. Visualization of the genome was performed using the CGView server (Stothard and Wishart, 2005), with CRISPRCasFinder (Couvin et al., 2018) and CARD (Alcock et al., 2019) modules for the visualization of CRISPR-Cas associated elements, and antibiotic resistance genes, respectively. This Whole Genome Shotgun project has been deposited at DDBJ/ENA/GenBank under the accession JAPKVL000000000. The version described in this paper is JAPKVL010000000.

### 2.3 BAC minimum inhibitory concentration (MIC) and doses-response curve determination

A bacterial inoculum of 10^6^ colony-forming units per milliliter (CFU/mL) was used to establish the minimum inhibitory concentration (MIC). MIC was determined with the broth microdilution method (Andrews, 2001) with slight modifications. A 200 µL was the total volume per well (96-well polypropylene microtiter plates, Greiner Bio One, Germany), and BAC was assessed after 36 hours at 30 °C (plate reader Biotek Epoch 2). The MIC was defined as the lowest concentration inhibiting liquid culture growth. The BAC (Fluka® Analytical, Germany) concentrations tested were: 0.7 µg/mL, 7.0 µg/mL, 14.0 µg/mL, 19.0 µg/mL, 23.0 µg/mL, 28.0 µg/mL, 33.0 µg/mL, 38.0 µg/mL, 76.0 µg/mL, and 380.0 µg/mL. For each concentration, seven replicates were tested. Additionally, as control, one well was used to blank for BAC + LB medium turbidity, seven wheels were used as growing control (without BAC), and seven wheels were used as LB medium turbidity control. Additionally, as control, one well was used to blank for BAC + LB medium turbidity, seven wells were used as growing control (without BAC), and seven wells were used as LB medium turbidity control. Optical density measurements in the plate reader were collected with the software Gen5 3.09. Growth curve analysis used the Growthcurver package for R (Sprouffske and Wagner, 2016). For analyzing differences in growing parameters, a Kruskal-Wallis test was conducted with software R version 4.1.1 (2021-08-10) (R Development Core Team, 2015). For the dose-response curve, a growth rate was calculated from OD_600_ measurement, taken every 10 minutes interval in LB medium (with and without BAC), using an algorithm described by Swain (Swain et al., 2016) and implemented in Python 3.6. Finally, the time-kill curve to BAC was determined by exposing exponential phase bacteria at a density of ~ 1×10^9^ CFU/mL to 40.0 µg/mL (Data non shown).

### 2.4 Antibiotic susceptibility profile

Approximately 10^6^ CFU /mL were inoculated onto semisolid LB Lennox Agar plates (LB medium, Carl Roth, Germany; plus agar 2%), following the disk diffusion method previously described (CLSI, 2018). Filter disks were impregned with 10 µL of the following antibiotics: ciprofloxacin (1 mg/mL), streptomycin (50 µg/mL), ceftazidime (50 µg/mL), trimethoprim (30 µg/mL), fosfomycin (30 µg/mL), colistin (20 µg/mL), daptomycin (4 µg/mL), rifampicin (62 µg/mL), gentamicin (8 µg/mL), chloramphenicol (30 µg/mL), ampicillin (100 µg/mL), doxycycline (100 mg/mL), amoxicillin (2048 µg/mL), vancomycin (20 mg/mL), and tetracycline (15 mg/mL). Additionally, a MIC estimation was established using E-test strips (BioMérieux®, France) for the following antibiotics: ampicillin, ciprofloxacin, fosfomycin, gentamicin, streptomycin, tobramycin, and ceftazidime.

### 2.5 Swimming test

An overnight culture was inoculated with a sterile toothpick on swimming plates. The swimming motility plates were prepared with 0.3% agar, 1.0% peptone (Acros Organics, Germany), and 0.5% NaCl (≥99.5% Carl Roth, Germany) (Zorzano et al., 2005).

### 2.6 Biofilm production test

The assay was done in 96-well microtiter dishes made of polypropylene (Greiner Bio One, Germany) as was previously described (Di Martino et al. 2005). Bacterial cells of overnight cultures at 30 °C were adjusted to 10^7^ UFC/mL in LB Lennox Medium (Carl Roth, Germany). Plates were inoculated with the bacterial suspensions (100 µl per well) and incubated at 30 °C for 24 h, 48 h, and 72 h. Next, the top of the plate was washed with water and placed in a new plate with 100 µl of crystal violet (1%) (O’Toole and Kolter, 1998). The plates were incubated for 30 min at room temperature and rinsed thoroughly and repeatedly with water. Finally, the dye was solubilized in acetic acid 35% (100 µl per well). The absorbance was determined at 595 nm. Each result represents 40 independent replicates of the experiment. For analyzing differences in biofilm formation, a Kruskal-Wallis test was conducted with software R version 4.1.1 (2021-08-10) (R Development Core Team, 2015).

### 2.7 Sample preparation and liquid chromatography–label-free quantification mass spectrometry (LC–LFQ-MS)

*Aeromonas hydrophila* proteomic analysis was carried out on exponential growing cultures treated with different BAC concentrations (9.5 µg/mL, 19 µg/mL, 38 µg/mL) for 30 minutes at 30 °C. Non-treated samples were used as control. Each experiment consisted of six replicates. After the exposition period, bacterial cells were pelleted by centrifugation at 4 °C, 10 000 x g for 2 minutes, the supernatant was discharged, and the pellet was kept at −80 °C. The bacterial pellets were treated with 100 µL of urea denaturing buffer (6 M urea, 2 M thiourea, and 10 mM HEPES, pH 8.0) followed by 8 cycles of freezing-unfreezing. All further sample preparation steps were performed as previously published (Rodríguez-Rojas and Rolff, 2020). After the digestion of proteins into peptides, salts, and contaminations (e.g. BAC) were removed before LC-MS analysis by SDB-RPS StageTips (Empore™ 2241) (Rappsilber et al., 2007). After elution from the StageTips, peptides were dried under vacuum.

Peptides were reconstituted in 30 μL of 0.05% TFA, 2% acetonitrile in water, and 1 μL were analysed by a reversed-phase capillary nano liquid chromatography system (Ultimate 3000, Thermo Scientific) connected to a Q Exactive HF mass spectrometer (Thermo Scientific). Samples were injected and concentrated on a trap column (PepMap100 C18, 3 μm, 100 Å, 75 μm i.d.×2 cm, Thermo Scientific) equilibrated with 0.05% trifluoroacetic acid in water. After switching the trap column inline, LC separations were performed on a capillary column (Acclaim PepMap100 C18, 2 μm, 100 Å, 75 μm i.d.×25 cm, Thermo Scientific) at an eluent flow rate of 300 nL/min. Mobile phase A contained 0.1% formic acid in water, and mobile phase B contained 0.1% formic acid in 80% acetonitrile / 20% water. The column was pre-equilibrated with 5% mobile phase B, and peptides were separated using a gradient of 5–44% mobile phase B within 35 min. Mass spectra were acquired in a data-dependent mode utilizing a single MS survey scan (m/z 300–1650) with a resolution of 60,000 in the Orbitrap, and MS/MS scans of the 15 most intense precursor ions with a resolution of 15,000. HCD fragmentation was performed for all peptide ions with charge states of 2+ to 5+ using a normalized collision energy of 27 and an isolation window of 1.4 m/z. The dynamic exclusion time was set to 20 s. Automatic gain control (AGC) was set to 3×10^6^ for MS scans using a maximum injection time of 20 ms. For MS2 scans, the AGC target was set to 1×10^5^ with a full injection time of 25 ms.

### 2.8 LC-MS data processing and protein quantification

MS and MS/MS raw data were analyzed using the MaxQuant software package (version 2.0.3.0) with the implemented Andromeda peptide search engine (Tyanova et al., 2016a). For the database search, a FASTA formatted protein sequence database of *A. hydrophila* downloaded from Uniprot (Bateman et al., 2023) (4,121 proteins, taxonomy 380703, March 22, 2022) was used. Filtering and statistical analysis was carried out using the software Perseus software version 1.6.14 (Tyanova et al., 2016b). Only proteins which were identified with LFQ intensity values in at least three out of 6 replicates (within at least one of the 4 experimental groups) were used for downstream analysis. Missing values were replaced from a normal distribution (imputation), using the default settings (width 0.3, downshift 1.8). Mean log2-fold differences between BAC-treated vs. non-treated control were calculated in Perseus, using Student’s t-tests with permutation-based FDR of 0.05. Proteins are defined as significantly regulated if they express a q-value < 0.05 (corresponds to the adjusted p-value) and at least a 2-fold change in intensity. The increased or decreased degree of overlapping proteins were represented as Venn diagrams using the online tool Venny (Oliveros, n.d.). Finally, the ontology and network of the identified proteins were retrieved with Uniprot database (Bateman et al., 2023) and STRING database (Szklarczyk et al., 2019) to obtain its functions and relations.

## 3 Results and Discussion

First, from an activated sludge previously exposed to BAC 10 mg/L (Chacón et al., 2021) a bacterial inoculum was transferred to TSA plates enriched to 25 mg/mL of BAC. After 24 h incubation at 28 °C, only less susceptible bacteria formed colonies. DNA extraction and *16S rRNA* gene sequencing was conducted in Macrogen® (South Korea). Of the 20 different identified isolates, 8 were identified as *Aeromonas hydrophila* with 99% of identity. Preliminary tolerance test showed that INISA09 presented a low susceptibility to BAC. Thereafter, we sequenced the bacterial genome of *A. hydrophila* INISA09 (SAMN31559380). The draft genome had an approximate size of 4.6 Mbp, 61.3% GC content, 121 tRNA genes, 14 rRNAs, 1 tmRNA, 50 miscRNAs, and 4,087 CDS. There were no CRISPR sequences or prophages present in the genome. These characteristics are in common with the reference strain *A. hydrophila* ATCC 7966 (Seshadri et al., 2006). The draft genome contained 4,273 genes. In Figure 1, a visualization of the genome is presented, including CRISPR-Cas associated proteins and antibiotic resistance genes; additionally an ANI comparison between INISA09 and ATCC 7966 *A. hydrophila* strain is shown; in general terms, both strains present a conservated cluster of genes; however, some arrangements in its genome position is observed. An analysis of single nucleotide polymorphisms (SNPs) showed 106,503 mutations (compared to *A. hydrophila* ATCC 7966), with 16,099 changes classified as non-synonymous mutations and 15,762 corresponding to missense variants. These mutations were found in genes associated with transport, antimicrobial resistance, gene expression regulation, motility, and redox activity. The strain *A. hydrophila* INISA09 underwent a profound genome rearrangement and genome plasticity compared to other strains, probably fine-tuned by thousands of mutations for better survival in the heavily contaminated environment. The number of changes is so huge that it is impossible to tackle each genomic change’s significance using the classic microbial genetic approach.

**Figure 1.**
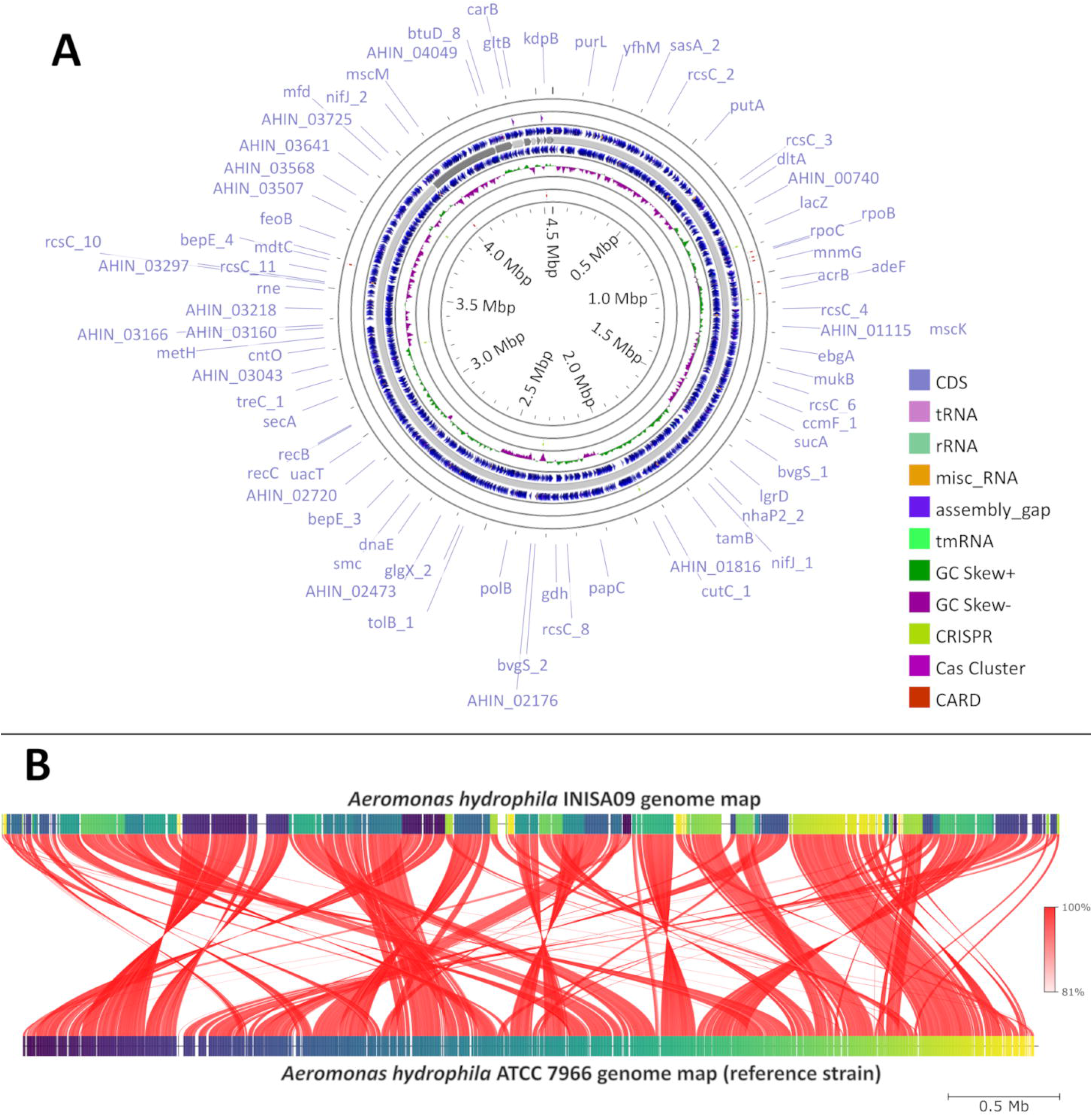
*Aeromonas hydrophila* INISA09 genome description. A. Visualization of the genome performed using the CGView server (Stothard and Wishart, 2005), with CRISPRCasFinder (Couvin et al., 2018)and CARD (Alcock et al., 2019) modules for the visualization of CRISPR-Cas associated protein, and antibiotic resistance genes, respectively. B. Average Nucleotide Identity (ANI) comparison between *Aeromonas hydrophila* INISA09 and *A. hydrophila* ATCC 7966. The red lines represent the reciprocal mapping among both strains indicating the conservation degree.

While exploring in the genome, focusing on genes with known roles in antimicrobial resistance, we found mutations in 112 loci associated with transport functions, 33 related to antimicrobial resistance, and 33 related to integral membrane composition. In addition, we identified mutations in genes such as *bcr* (bicyclomycin resistance protein), *arnA* (bifunctional polymyxin resistance protein ArnA), *drrA* (daunorubicin/doxorubicin resistance ATP-binding protein DrrA), *fsr* (fosmidomycin resistance protein), *emrB, mdtE, mdtA, mdtH, mdtL, norM, mdlB* (all multidrug resistance proteins), and *qacC* (quaternary ammonium compound-resistance protein QacC). The mutations in genes associated with efflux pumps include *acrB* (efflux pump AcrB), *acrA* (efflux pump AcrA), *acrE* (efflux pump AcrE), and the outer membrane protein *tolC* (outer membrane protein), which is part of the AcrAB-TolC multidrug efflux pump complex (Tikhonova and Zgurskaya, 2004). These results are consistent with a previous study conducted on *Salmonella enterica* serovar Typhimurium showing that during exposure to BAC, TolC protein connects with AcrAB efflux pump system to expel the biocide to the extracellular space (Guo et al., 2014).

We also identified intragenic mutations in genes that code for membrane-related proteins such as the porins *ompA* (outer membrane protein A) and *ompW*□ (outer membrane protein W), both also related to the adhesion process and biofilm formation (Maiti et al., 2012; Wang et al., 2022). The porins *ompD* (outer membrane protein D) and *ompN* (outer membrane protein N) also showed mutations in the INISA09 strain, which might confer low susceptibility to benzalkonium chloride (BAC) or other biocides since cationic surfactants target the outer membrane as the primary site (Gilbert and Moore, 2005). In a previous study in *Escherichia coli*, some of these genes were differentially expressed under BAC exposure in a transcriptomic assay (Forbes et al., 2019). In an *in vitro* experiment in *E. coli*, BAC induced tolerance mediated by the genes *lpxM*. This gene codes for a myristoyl transferase responsible for the last acylation step in lipid A synthesis (Nordholt et al., 2021). The wild *A. hydrophila* INISA09 genome presented 18 mutations in this gene; three were non-synonymous mutations: Y21H, M59I, and Y225H. However, these mutations differ from those observed in *E. coli:* A96E, M148R, L207I, A242E, and W298G.

We analyzed the antibiotic susceptibility of *A. hydrophila* INISA09 strain by E-test and disk diffusion assays. The MIC estimated by E-test showed different degrees of susceptibility to seven antibiotics, including ampicillin >256 µg/mL, ciprofloxacin 0.064 µg/mL, fosfomycin 8 µg/mL, gentamycin 1 µg/mL, streptomycin 8 µg/mL, tobramycin 1.5 µg/mL and ceftazidime 0.032 µg/mL. A previous study conducted in environmental isolates of *Aeromonas* sp. (Goñi-Urriza et al., 2000) showed high resistance to penicillins such as ampicillin and increased susceptibility to the third generation of cephalosporines. Our results were similar for the susceptibility of *A. hydrophila* INISA09 to the third generation of cephalosporines, indicating the absence of extended-spectrum beta-lactamases. The values obtained when compared to those of Goñi *et al*.(Goñi-Urriza et al., 2000) were the same for gentamycin (1 µg/mL), lower for streptomycin (16 µg/mL) and third-generation cephalosporins (cefotaxime 8 µg/mL). We highlight that strain INISA09 was highly susceptible to fosfomycin (0.2-8 µg/mL).

The European Committee on Antimicrobial Susceptibility Testing (EUCAST) (EUCAST, 2022) informed that all clinical isolates of□*A. hydrophila*□display intrinsic resistance to ampicillin, amoxicillin, amoxicillin-clavulanic acid, ampicillin-sulbactam, and cefoxitin. *A. hydrophila* is intrinsically resistant to benzylpenicillin, glycopeptides, lipoglycopeptides, fusidic acid, lincosamides, streptogramins, rifampicin, oxazolidines, and macrolides (except azithromycin). In this case, *A. hydrophila* INISA09 was only resistant to ampicillin (penicillin) and susceptible to ciprofloxacin (quinolone), tobramycin, streptomycin, and gentamycin (aminoglycosides), ceftazidime (cephalosporine), and fosfomycin (epoxide). It is important to mention that European Centre for Disease Prevention and Control (ECDC) criteria define multi-drug resistance (MDR) as acquired non-susceptibility to at leat one agent in three or more antimicrobial categories (Magiorakos et al., 2012). Table 1 details the antibiotics tested and their phenotype according to the recommendations of EUCAST for *Aeromonas sp*. and *Vibrio sp*. (another Gammaproteobacteria, closely phylogenetically related), and in Table S1, we show the tested concentration and inhibition halo diameter.

**Table 1.** Antibiotic susceptibility testing of *Aeromonas hydrophila* INISA09 determined by disk diffusion method for different antibiotic families. See Supplementary Table 1 for comple information such as inhibition hallo diametes and tested concentrations.

| Therapeutic class | Antimicrobial | Phenotype |
| --- | --- | --- |
| Aminoglycoside | Gentamicin | Susceptible |
|  | Streptomycin | Susceptible* |
| Cephalosporine | Ceftazidime | Susceptible* |
| Dihydropyrimidine | Trimethoprim | Susceptible* |
| Epoxide | Fosfomycin | Susceptible |
| Glycopeptide | Vancomycin* | Intrinsic resistant |
| Lipopeptide | Daptomycin* | Intrinsic resistant |
| Penicillin | Amoxicillin* | Intrinsic resistant |
|  | Ampicillin* | Intrinsic resistant |
| Phenicol | Chloramphenicol | Susceptible |
| Polymyxin | Colistin | Susceptible |
| Quinolone | Ciprofloxacin | Sensible |
| Rifamycin | Rifampicin* | Intrinsic resistant |
| Tetracycline | Doxycycline | Susceptible * |
|  | Tetracycline | Susceptible* |

\* *Vibrio* sp. breakpoint according to EUCAST (EUCAST, 2022).

We analyzed the effect of different near-MIC BAC concentrations on the growth curve of the INISA09 strain. For this, we used the Grothcurver R package, which fits growth curve data to the standard form of the logistic equation commonly used in ecology and evolution (Sprouffske and Wagner, 2016); the parameters estimates are shown in Figure S1. To this end, we employed the estimated MIC for BAC (38.0 µg/mL) to evaluate parameters related to growth, such as the growth rate (*r*-value), the maximum population size or carrying capacity (*K*-value), the generation time, and the empirical area under the curve (auc_e_) estimated from the obtained optical density (OD^600^) data. We compared these values with a non-treated control using the R package Growthcurver (Sprouffske and Wagner, 2016).

In Table 2, we present the results for relevant growth curve parameters (see Supplementary Figure 1). We found significant statistical differences for the carrying capacity (*K*, which describes the maximum reachable biomass of the culture, *p=*1.5 × 10^−7^) and the empirical area under the curve parameters (auc_e_, another global parameter that integrates other parameters such as including the initial population size, maximum growth rate, carrying capacity etc, *p=*7.7 × 10^−7^). In a dose-response curve analysis, the EC_50_ (half maximal effective concentration of BAC in 30 minutes) for INISA09 corresponded to 33.0 µg/mL. Figure 2 shows a typical growth curve behavior of *A. hydrophila* under three different BAC concentrations: 9.5 µg/mL, 19 µg/mL and 38 µg/mL, and the average area under the curve obtained. This strain’s resistance level is notably higher than other bacteria such as *Escherichia coli*, where a tolerant population presents a MIC of 12 µg/mL (Moen et al., 2012).

**Table 2.** Growing descriptive parameters obtained for INISA09 strain during a 38.0 µg/mL BAC exposition and non-BAC exposure. See also Supplementary Figure 2.

| Parameter | BAC exposed | Non-BAC exposed |
| --- | --- | --- |
| *r*-value | 1.8 x 10<sup>-3</sup> | 6.2 x 10<sup>-3</sup> |
| *K*-value* | 1.5 x 10<sup>-3</sup> | 1.1 |
| auc<sub>e</sub>* | 1.7 | 968 |
| Generation time | 877 minutes | 111 minutes |

\* Statistical difference $p \leq 0.05$

**Figure 2.**
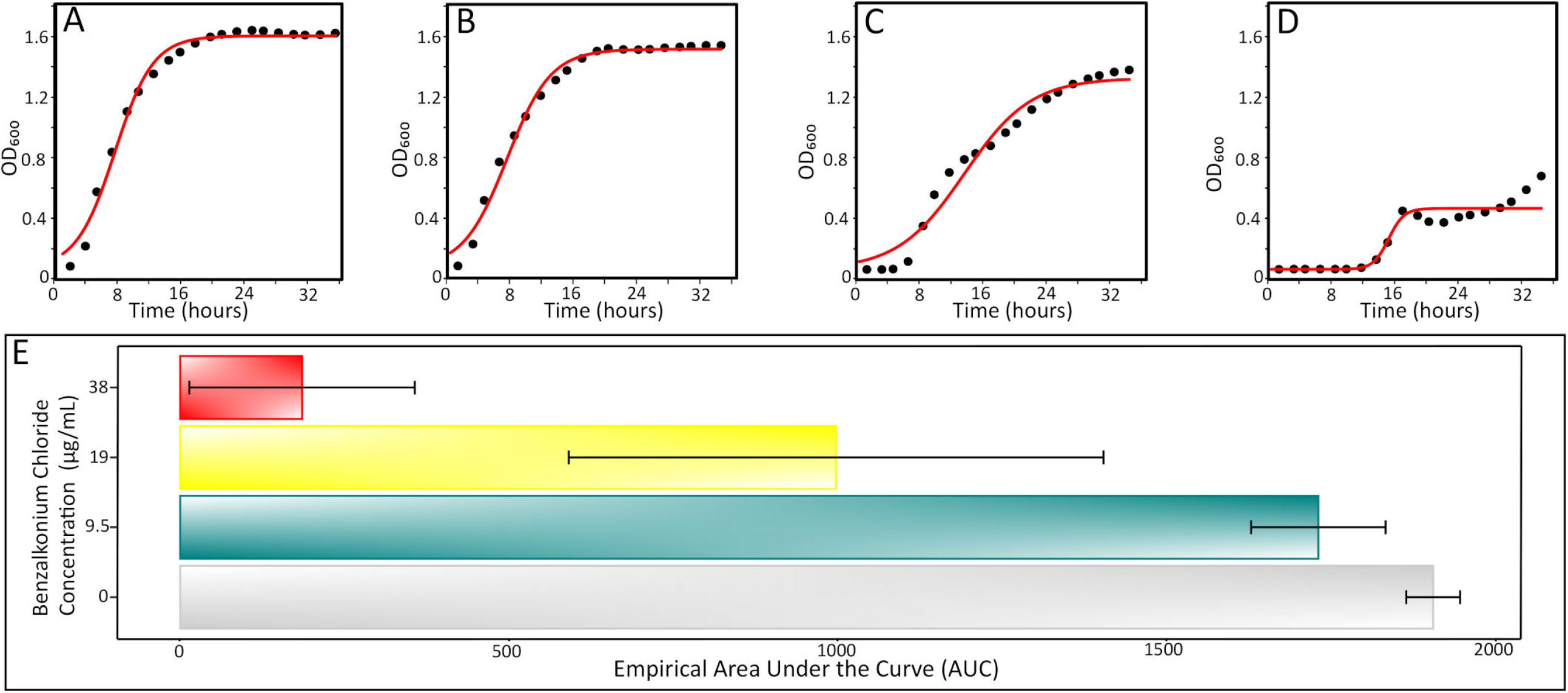
Typical growth curves of *A. hydrophila* INISA09 exposed to different BAC concentrations. A: Non-treated control, B: BAC 9.5 µg/mL, C: BAC 19 µg/mL, D: 38 µg/mL doses; and E: empirical area under the curve, a global parameter that integrates other arametes such as growth rate, carriying capacity, death phase, initial population size, etc. The red line represents the logistic regression model obtained from the absorbance data and the empirical AUC was obtained from the experimental absorbance obtained curve from a five independent repetiions according to previously described procedure (Sprouffske and Wagner, 2016).

We also tested bacterial mobility using the swimming motility plate assay (Zorzano et al., 2005). *A. hydrophila* INISA09 showed a moderate swimming capacity. Additionally, a biofilm formation assay shows a strong increase in biofilm mass in the first 48 hours (Figure S2). The statistical analysis showed differences in biofilm production in the three tested periods (*p*= 1.1 × 10^−3^). However, post hoc analyses indicated that the differences occur among the first 48 hours (*p*= 0.03), whereas the difference between 48 and 72 hours is not significant (*p*= 0.16). This result indicates that *A. hydrophila* biofilms mature within 48 h under experimental conditions.

Next, we performed a label-free quantification LC-mass spectrometry proteomic analysis to understand the decreased susceptibility to BAC of strain *A. hydrophila* INISA09. To this end, we exposed the cells for 30 min to different concentrations of BAC and compared them to the non-treated control. The chosen BAC concentrations were 9.5, 19, and 38 µg/mL, corresponding to ¼, ½, and the MIC but the experiments were carried out with a higher inoculm (aproximaterly 10^8^ CFU/mL) to have more material for the protein extraction. We found that 111, 379, and 233 proteins with a significant change in relative abundance when the bacteria were treated with ¼ MIC (9.5 µg/mL), ½ MIC (19 µg/mL), and the MIC (38 µg/mL), respectively (Figure 3). Volcano plots representing the differential expression of all quantified proteins are shown in Figure 4. Tables 3 and 4 lists 15 proteins with the largest fold change in each experiment. The whole dataset is shown in Supplementary Table 3.

**Figure 3.**
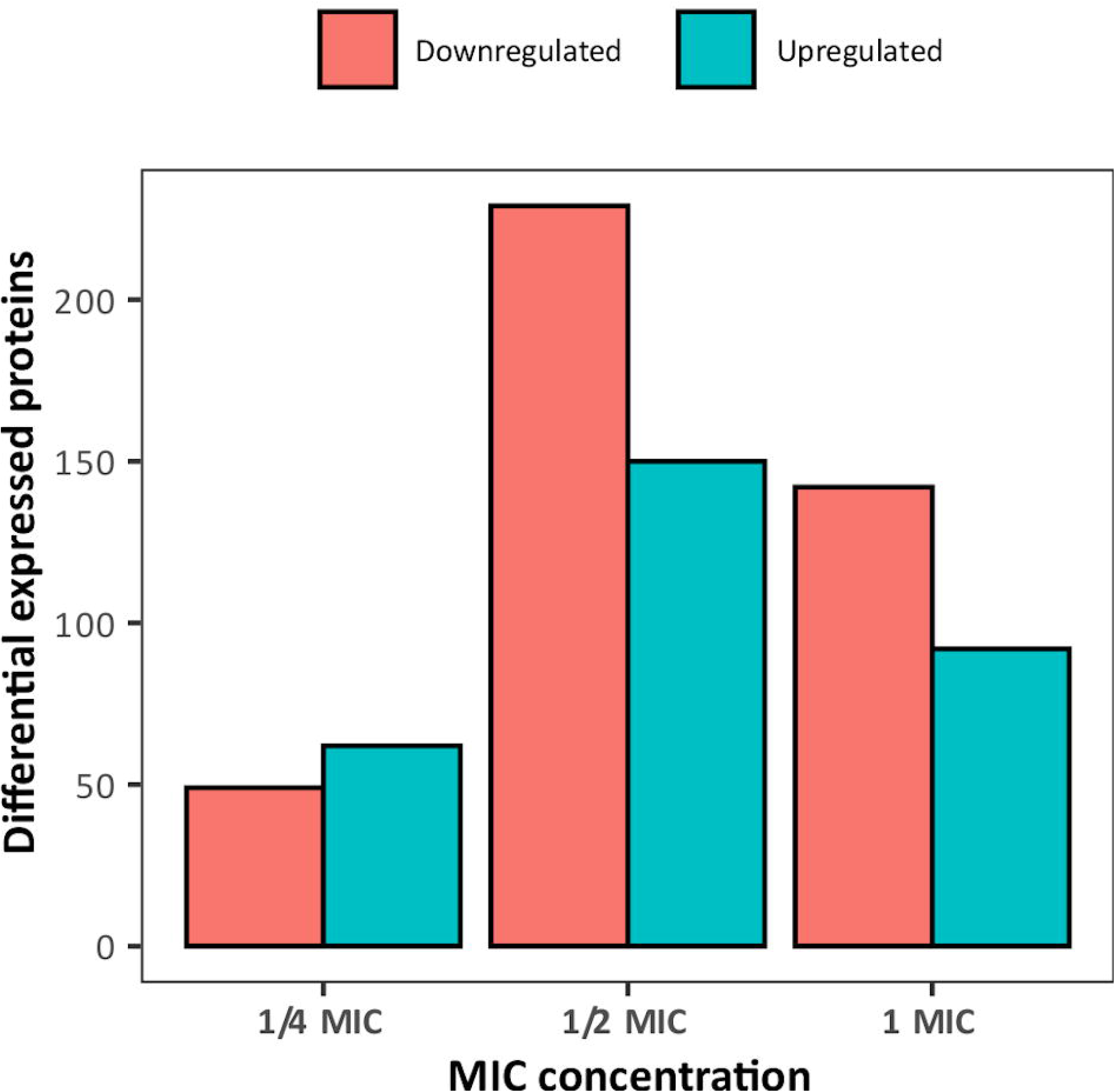
Overview of the number of differentially expressed proteins in *Aeromonas hydrophila* INSA09 after 30 minutes exposure to BAC (¼ MIC: 9.5 µg/mL, ½ MIC: 19 µg/mL, and MIC: 38 µg/mL doses).

**Figure 4.**
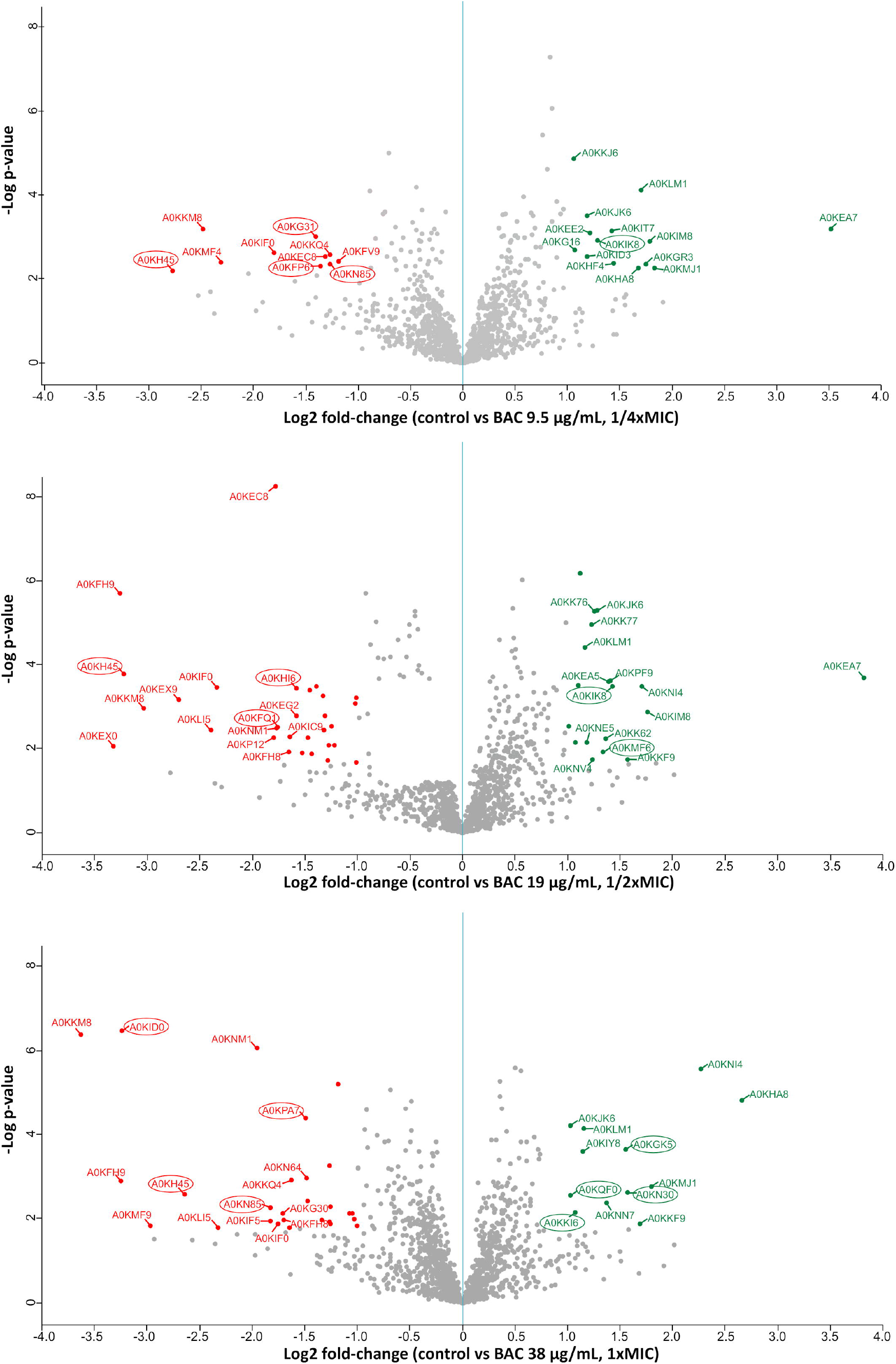
Label-free quantification LC-MS proteomics results. Volcano plot showing the log2 fold-change of mean relative protein intensities of *A. hydrophila* determined by LC-MS and label-free quantification (BAC-treated vs. non-treated control). Proteins with significant higher abundance in non-treated controls are coloured in red (downregulated upon BAC-treatment), while proteins with significant higher abundance after BAC-treatment are coloured in green (upregulated). Proteins are defined as significantly regulated if they express a q-value < 0.05 (FDR adjusted p-value) and at least a 2-fold change in intensity. Proteins marked with a circle correspond to membrane-related proteins, according to the Uniprot database (Bateman et al., 2023).

**Table 3.**
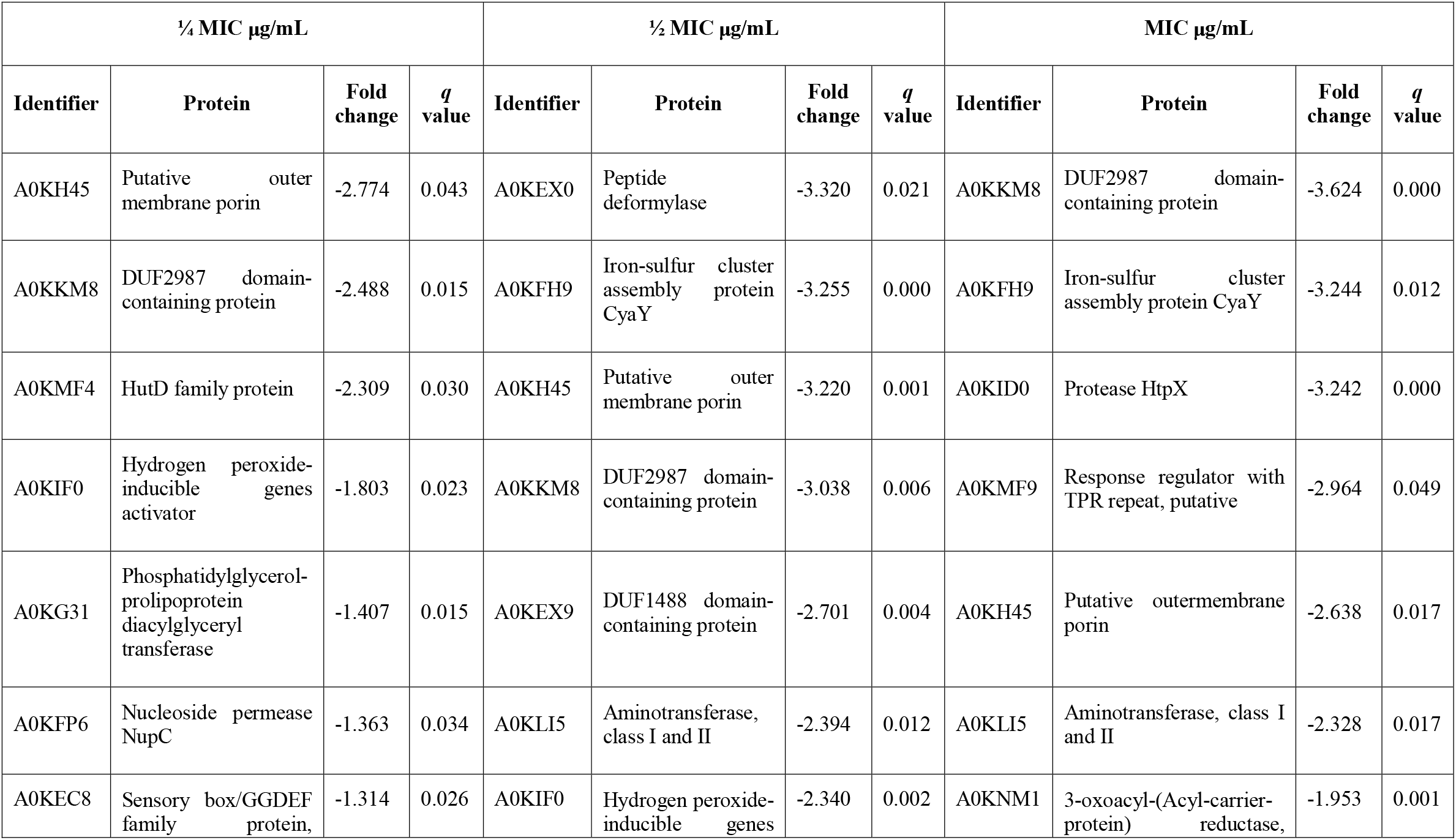
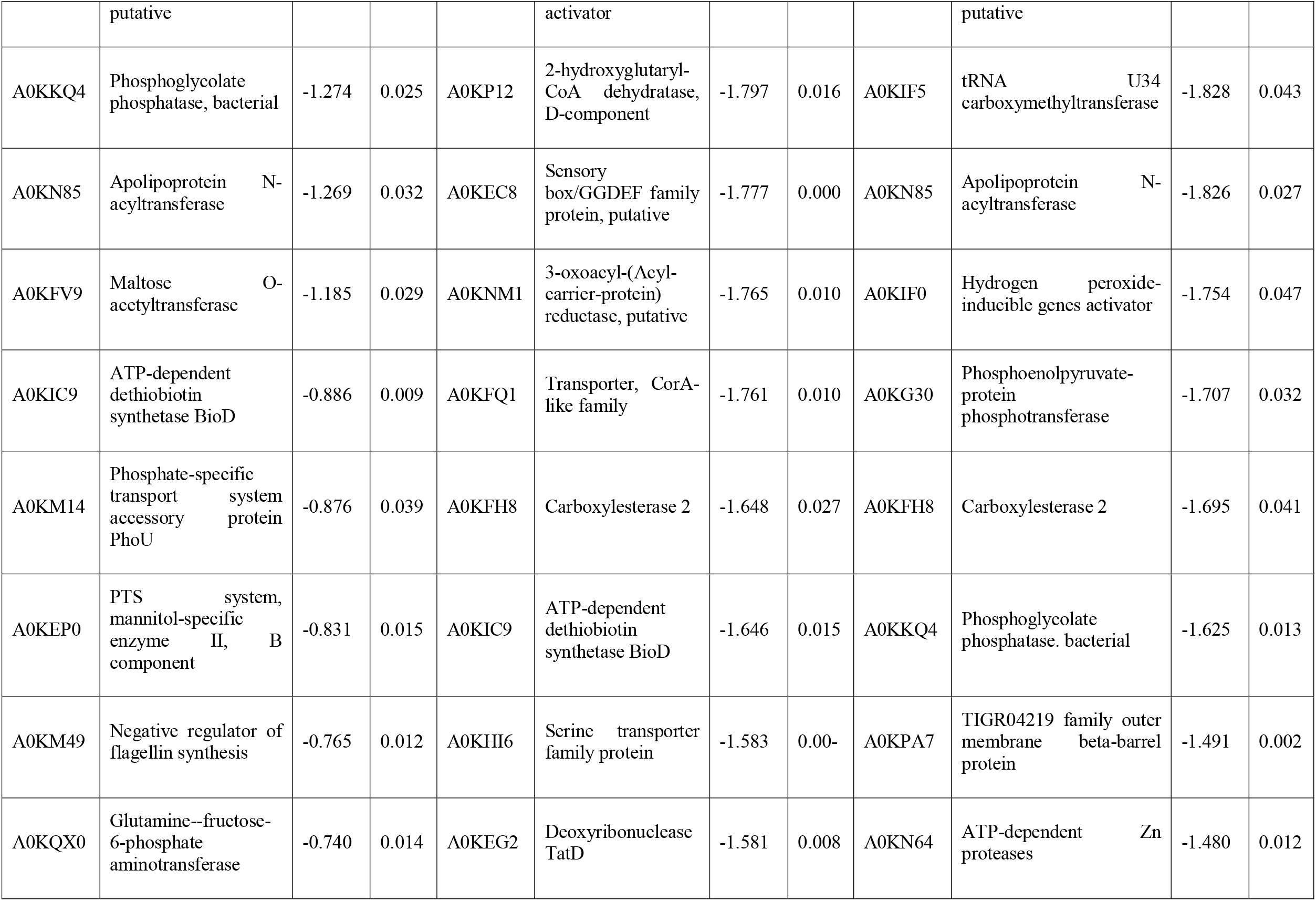
Downregulated proteins for at least a 2-fold change in intensity, measured by LC-MS using label-free quantification, in *A. hydrophila* after been exposed to three different BAC concentrations. q value < 0.05 corresponds to adjusted p values after false discovery rate control (FDR).

| ¼ MIC µg/mL |  |  |  | ½ MIC µg/mL |  |  |  | MIC µg/mL |  |  |  |
| --- | --- | --- | --- | --- | --- | --- | --- | --- | --- | --- | --- |
| Identifier | Protein | Fold change | q value | Identifier | Protein | Fold change | q value | Identifier | Protein | Fold change | q value |
| A0KH45 | Putative outer membrane porin | -2.774 | 0.043 | A0KEX0 | Peptide deformylase | -3.320 | 0.021 | A0KKM8 | DUF2987 domain-containing protein | -3.624 | 0.000 |
| A0KKM8 | DUF2987 domain-containing protein | -2.488 | 0.015 | A0KFH9 | Iron-sulfur cluster assembly protein CyaY | -3.255 | 0.000 | A0KFH9 | Iron-sulfur cluster assembly protein CyaY | -3.244 | 0.012 |
| A0KMF4 | HutD family protein | -2.309 | 0.030 | A0KH45 | Putative outer membrane porin | -3.220 | 0.001 | A0KID0 | Protease HtpX | -3.242 | 0.000 |
| A0KIF0 | Hydrogen peroxide-inducible genes activator | -1.803 | 0.023 | A0KKM8 | DUF2987 domain-containing protein | -3.038 | 0.006 | A0KMF9 | Response regulator with TPR repeat, putative | -2.964 | 0.049 |
| A0KG31 | Phosphatidylglycerol-prolipoprotein diacylglycerol transferase | -1.407 | 0.015 | A0KEX9 | DUF1488 domain-containing protein | -2.701 | 0.004 | A0KH45 | Putative outermembrane porin | -2.638 | 0.017 |
| A0KFP6 | Nucleoside permease NupC | -1.363 | 0.034 | A0KLI5 | Aminotransferase, class I and II | -2.394 | 0.012 | A0KLI5 | Aminotransferase, class I and II | -2.328 | 0.017 |
| A0KEC8 | Sensory box/GGDEF family protein, | -1.314 | 0.026 | A0KIF0 | Hydrogen peroxide-inducible genes | -2.340 | 0.002 | A0KNM1 | 3-oxoacyl-(Acyl-carrier-protein) reductase, | -1.953 | 0.001 |
|  | putative |  |  |  | activator |  |  |  | putative |  |  |
| A0KKQ4 | Phosphoglycolate phosphatase, bacterial | -1.274 | 0.025 | A0KP12 | 2-hydroxyglutaryl-CoA dehydratase, D-component | -1.797 | 0.016 | A0KIF5 | tRNA U34 carboxymethyltransferase | -1.828 | 0.043 |
| A0KN85 | Apolipoprotein N-acyltransferase | -1.269 | 0.032 | A0KEC8 | Sensory box/GGDEF family protein, putative | -1.777 | 0.000 | A0KN85 | Apolipoprotein N-acyltransferase | -1.826 | 0.027 |
| A0KFV9 | Maltose O-acyltransferase | -1.185 | 0.029 | A0KNM1 | 3-oxoacyl-(Acyl-carrier-protein) reductase, putative | -1.765 | 0.010 | A0KIF0 | Hydrogen peroxide-inducible genes activator | -1.754 | 0.047 |
| A0KIC9 | ATP-dependent dethiobiotin synthetase BioD | -0.886 | 0.009 | A0KFQ1 | Transporter, CorA-like family | -1.761 | 0.010 | A0KG30 | Phosphoenolpyruvate-protein phosphotransferase | -1.707 | 0.032 |
| A0KM14 | Phosphate-specific transport system accessory protein PhoU | -0.876 | 0.039 | A0KFH8 | Carboxylesterase 2 | -1.648 | 0.027 | A0KFH8 | Carboxylesterase 2 | -1.695 | 0.041 |
| A0KEP0 | PTS system, mannitol-specific enzyme II, B component | -0.831 | 0.015 | A0KIC9 | ATP-dependent dethiobiotin synthetase BioD | -1.646 | 0.015 | A0KKQ4 | Phosphoglycolate phosphatase, bacterial | -1.625 | 0.013 |
| A0KM49 | Negative regulator of flagellin synthesis | -0.765 | 0.012 | A0KHI6 | Serine transporter family protein | -1.583 | 0.00- | A0KPA7 | TIGR04219 family outer membrane beta-barrel protein | -1.491 | 0.002 |
| A0KQX0 | Glutamine--fructose-6-phosphate aminotransferase | -0.740 | 0.014 | A0KEG2 | Deoxyribonuclease TatD | -1.581 | 0.008 | A0KN64 | ATP-dependent Zn proteases | -1.480 | 0.012 |

**Table 4.**
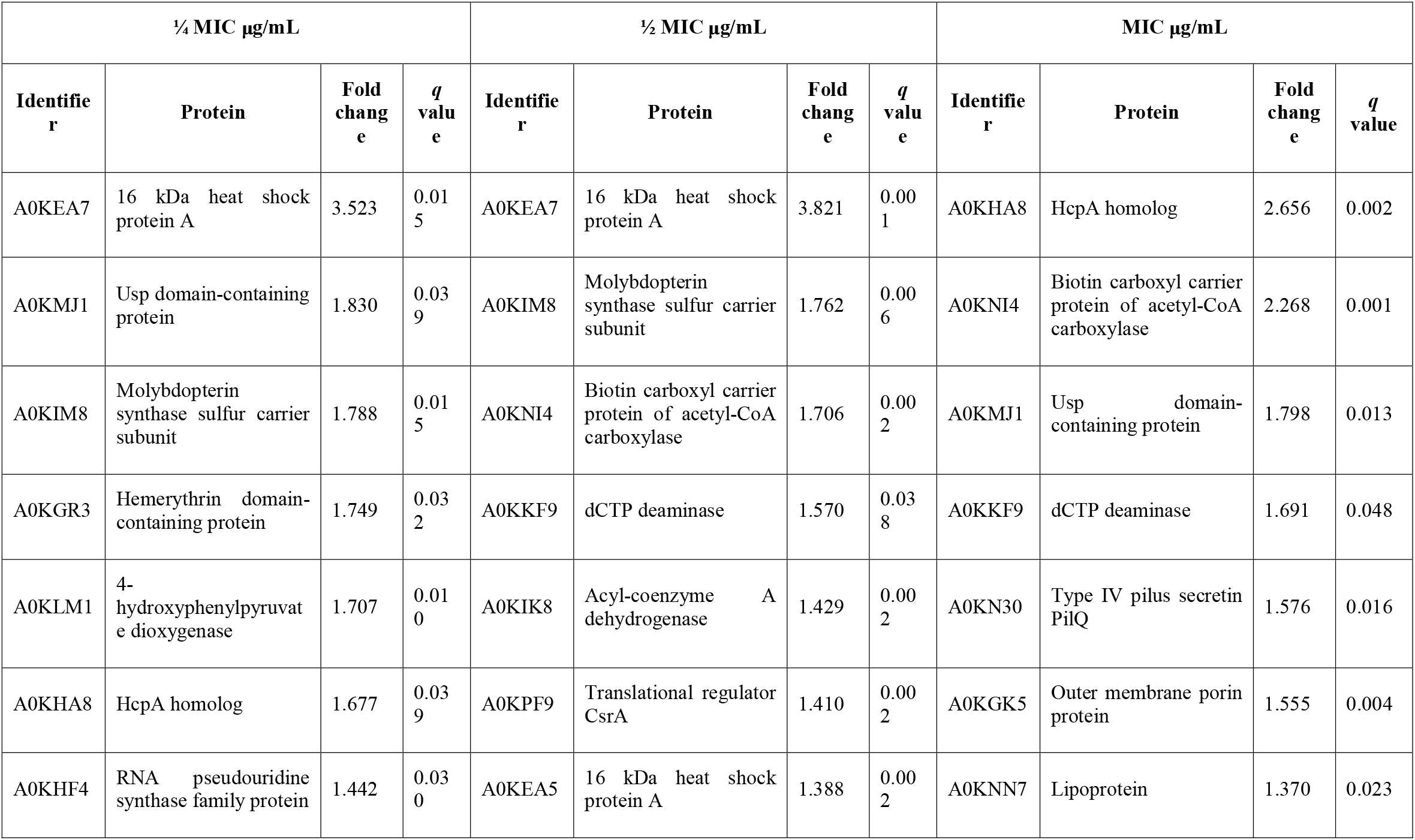
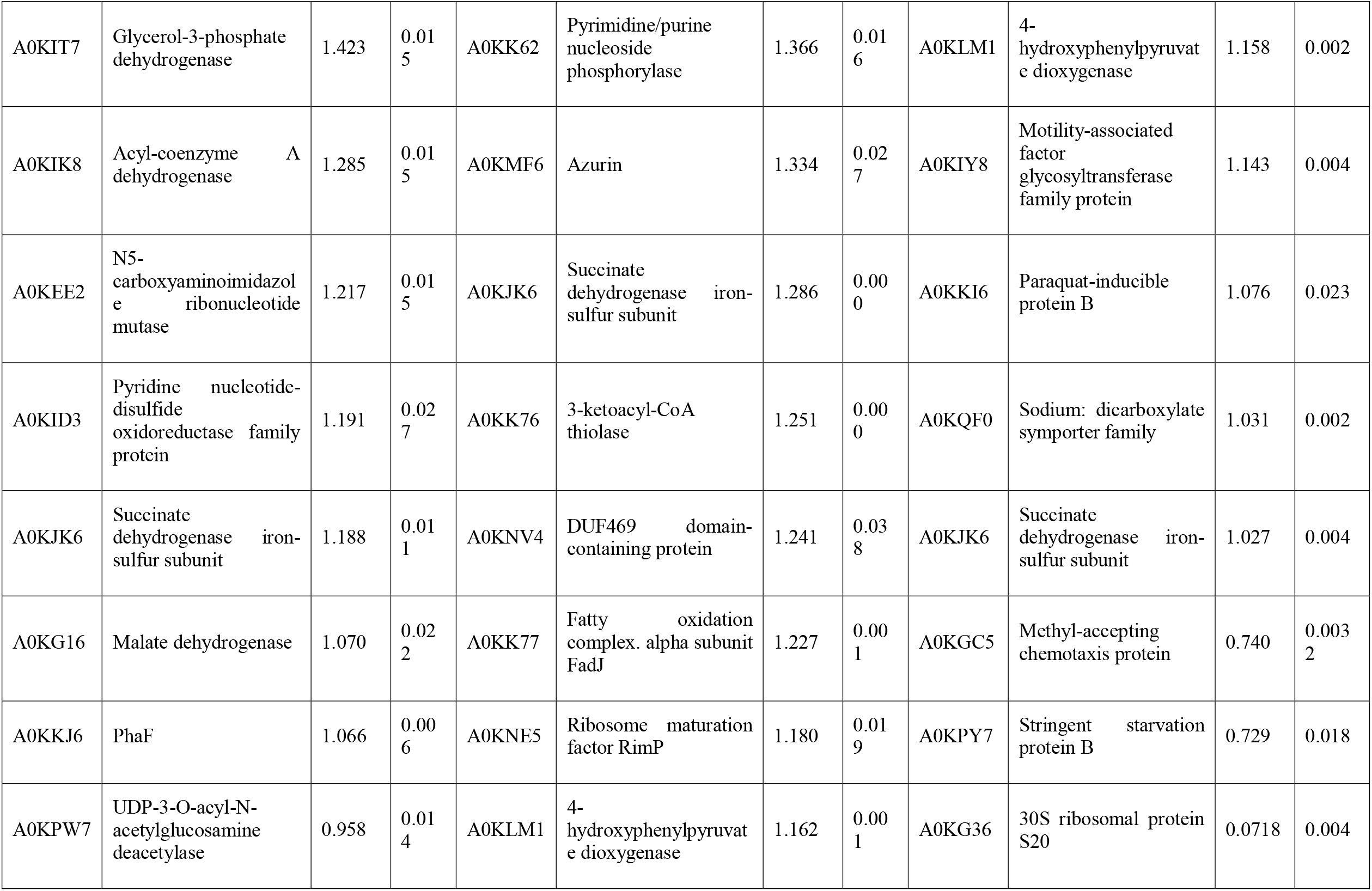
Upregulated proteins for at least a 2-fold change in intensity, measured by LC-MS using label-free quantification, in *A. hydrophila* after been exposed to three different BAC concentrations. q value < 0.05 corresponds to adjusted p values after false discovery rate control (FDR).

In Figure 5, we show a Venn diagram illustrating the common core set of protein expression and the disimirarities among the three used BAC concentrations. Twenty-one downregulated proteins upon BAC treatment consistently coincide across all three conditions. These include five envelope-related proteins, including the membrane proteins A0KH45, A0KHA5, and A0KI39, the cell wall degradation protein A0KHL2, and a carbohydrate-associated transporter A0KEP0. Two of these proteins are related to different stress responses: A0KIF0 is an activator of hydrogen peroxide-inducible genes, and A0KM49 is a negative regulator of flagellin synthesis. Another 12 expressed genes are responsible for metabolic pathways, such as nucleic acid and fatty acid metabolism. One is a disulfide-reductase enzyme (A0KEH0), and the other is an undetermined function protein (DUF-domain protein) (A0KKM8). In addition, the upregulated proteins include six enzymes associated with metabolic processes (A0KN18, A0KLM1, A0KP19, A0KPW7, and A0KQG0), two membrane-related proteins (A0KHC8 and A0KKM0, both with oxidoreductase activity associated with electron transport), and three proteins with binding functions (A0KKE2 to ribosomes, and A0KIA0 and A0KJK6 to ion reductase enzymes). The last one, A0KM63, is a DUF-domain protein with an unknown function.

**Figure 5.**
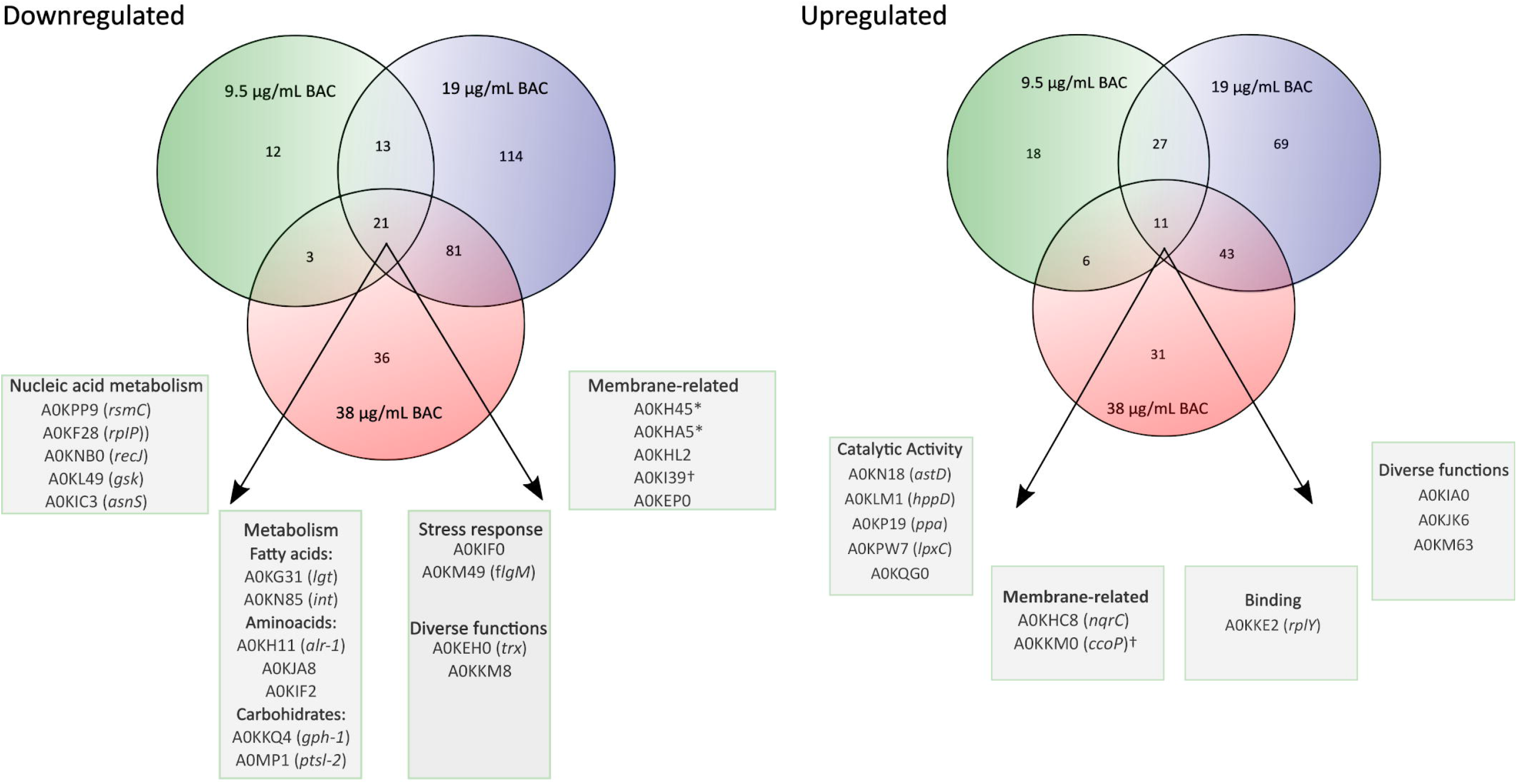
Venn diagrams show the overlap of proteins that show a significant change in their relative abundance (BAC-treated vs. non-treated control) using different BAC concentrations (¼ MIC −9.5 µg/mL-, ½ MIC-19 µg/mL-, and 1 MIC-38 µg/mL-). The porins are marked with * and the cytochromes are marked with †.

The results of the quantitative proteome analysis indicate that the basal response of *A. hydrophila* INISA09 to BAC exposure includes membrane rearrangements and stabilization of the oxidative cell status. Although there are few examples of environmental isolates exposed to BAC for comparative analysis, these results are consistent with those previously reported in a transcriptomic study of *Escherichia coli*. In this study, exposure to BAC reduced the expression of outer-membrane porins, decreased motility and chemotaxis, and increased expression of respiration-associated genes (Forbes et al., 2019).

A closer examination of the proteins with the highest differential expression level (Tables 3 and 4) reveals that three proteins are strongly downregulated across all three tested concentrations. The protein A0KH45 is described as a putative outer membrane porin and a member of the OprD family. This group of proteins is associated with the transport of essential amino acids, and mutations in its sequence can lead to carbapenem resistance (Bateman et al., 2023). A previous study reported that reduced levels of its expression in the outer membrane could allow carbapenem accumulation in outer membrane vesicles and reduce free carbapenems within the cell (Chevalier et al., 2017). Lowering the expression of porins, such as OmpC, OmpD, and OmpF, is one mechanism that decreases the susceptibility of Gram-negative bacteria to biocides (Ferenci and Phan, 2015). In this regard, the INISA09 strain presented mutations in ompD. It has been previously described that *A. hydrophila* downregulates porin expression under iron-restricted conditions (Wang et al., 2017). *A. hydrophila* may also use this mechanism to resist stress conditions, such as biocide exposure or starvation.

The protein A0KKM8 appears in the databases as an uncharacterized that contains the DUF2987 domain. According to the GenBank database, this domain is present in various *Vibrio* species, including *V. parahaemolyticus, V. nigripulchritudo, V. natriegens, V. cholerae, V. cyclitrophicus, V. antiquaries, V. lentus, V. alginolyticus, and V. mimicus*, as well as in other species such as *Pseudoalteromonas rubia, Alteromonas macleodii, Shewanella baltica*, and *Alteromonas mediterranea*. It is also present in several *Aeromonas* species, including *A. caviae, A. dhakensis, A. jandei, A. media, A. salmonicida, A. veronii, A. sobria*, and *A. enteropelogenes*. A protein-protein interaction analysis using the STRING database (Szklarczyk et al., 2019) reveals a co-occurrence with molecules such as BamC, a lipoprotein part of the Outer Membrane Protein Assembly Complex involved in the assembly and insertion of beta-barrel structures into the outer membrane, and some uncharacterized proteins. This protein has not been characterized in detail until now. However, it is likely a membrane-related protein common in aquatic bacteria and necessary for responding to stress. Therefore, we recommend studying the role of the protein A0KKM8 in future research via functional genomics to improve our understanding of its function, especially in biocide survival conditions.

The A0KIF0 protein, also known as Hydrogen Peroxide-Inducible Genes Activator, was downregulated in the presence of BAC. This protein has been described as part of the OxyR regulon since 1990, and its function is to defend the cell against oxidative damage caused by hydrogen peroxide (Storz et al., 1990). In *A. hydrophila*, the expression of OxyR can be modulated by the LahS transcriptional regulator. Under stress conditions, LahS diminishes OxyR expression and enhances biofilm formation (Dong et al., 2018). A similar regulation may occur in INISA09, which suggests that the stress response to BAC is not mediated by hydrogen peroxide-related genes.

Additionally, the protein A0KKM49 (FlgM) is also downregulated. This response under stress conditions has been observed previously in a *Salmonella* sp., where it was shown that damages in the flagella and membrane integrity could repress nutrient acquisition and activate the cell envelope stress response due to environmental factors affecting the bacterial envelop (Spöring et al., 2018). A similar mechanism is likely activated in *A. hydrophila* under BAC exposure.

For moderate and high doses of BAC, downregulation of several membrane proteins was observed, including A0KFH9 (iron-sulfur cluster assembly protein CyaY), which is related to the electron transport chain, radicals generation, and regulation of biosynthesis of sulfur derivatives (Iannuzzi et al., 2011), and transport proteins such as A0KFQ1 (a member of the CorA-like family of transporters with metal ion transporter activity), A0KID0 (with metalloendopeptidase activity and zinc ion binding), and A0KPA7 (an outer membrane protein). Besides, some proteins related to fatty acid syntheses, such as A0KN85, A0KFH8, and A0KNM1, were also downregulated, indicating bacterial membrane reorganization and an alternative response to oxidative stress as a consequence of BAC exposure.

In the INISA09 strain, several cytochrome-related and metal-binding proteins were upregulated during BAC treatment, including A0KIM8, A0KHC8, and A0KKM0. Cytochromes and respiratory chain proteins can stabilize the cell’s redox status without metal loss through an independent pathway of OxyR regulation (Mishra and Imlay, 2012). These results suggest that in response to BAC, membrane proteins, such as those involved in oxidative stress response, could be upregulated in the strain INISA09.

Under BAC stress conditions, several proteins related to fatty acid metabolism and membrane proteins were upregulated in the INISA09 strain. A0KK76 (FadI) and A0KK77 (FadJ) are two proteins regulated by the operon CO03404902 (Okuda and Yoshizawa, 2011), and both are part of the fatty acid oxidation with acetyl-CoA. According to STRING (Szklarczyk et al., 2019), these proteins interact with protein NuoC (A0KJ65), a membrane protein associated with electron transport across the membrane to conserve the redox energy in a proton gradient. Another upregulated protein observed that interacts with NuoC is A0KJK6 (the iron-sulfur subunit of succinate dehydrogenase), which is also regulated by the operon CO006649937 that regulates the expression of the succinate dehydrogenase-related genes (dhC, AHA_1924, sdhA, and AHA_1926). The overexpressed protein A0KNI4 (AccB) is part of an operon composed of the genes aroQ, accB, and accC, which are part of the acetyl coenzyme A carboxylase complex and involved in the fatty acid synthesis pathways.

Two other upregulated proteins, NifJ-1 and NifJ-2, are oxidoreductases that transfer electrons from pyruvate to flavodoxin and have iron centers. They interact directly with FadI, FadJ, AccB, A0KJK6, AstD (A0KN18), Mdh (A0KG16), and A0KIA0, all of which are related to oxidoreductase activities and fatty acid metabolism. This pattern is remarkable because most of the proteins mentioned have metal centers and can participate in electron transfer processes as mechanisms to balance oxidative stress disequilibrium (Crack et al., 2012). In the case of the INISA09 strain, the damage caused by BAC at the membrane level could be managed with a reorganization of the fatty acid composition since the primary redox reactions are related to fatty acid metabolism, using redox reactions to equilibrate the oxidative stress.

Finally, we found some correspondences between the most critically differentially expressed proteins (Tables 3 and 4) and the presence of missense SNPs (see Table 5) (stop codons, deletions, and insertions with frameshift changes were not found). The average number of missense SNPs is 3 per differentially expressed protein. However, it is remarkable that the downregulated proteins A0KH45 (porin), A0KHI6 (transporter), and A0KMF4 (HUT domain protein) present more than 10 missense SNPs. Regarding upregulated proteins, A0KKE3 (histidine kinase) and A0KNN7 (lipoprotein) present more than 15 missense mutations in their sequence. However, further research is required to determine the effect of these mutations on the global BAC response presented by INISA09 or other environmental bacteria.

**Table 5.**
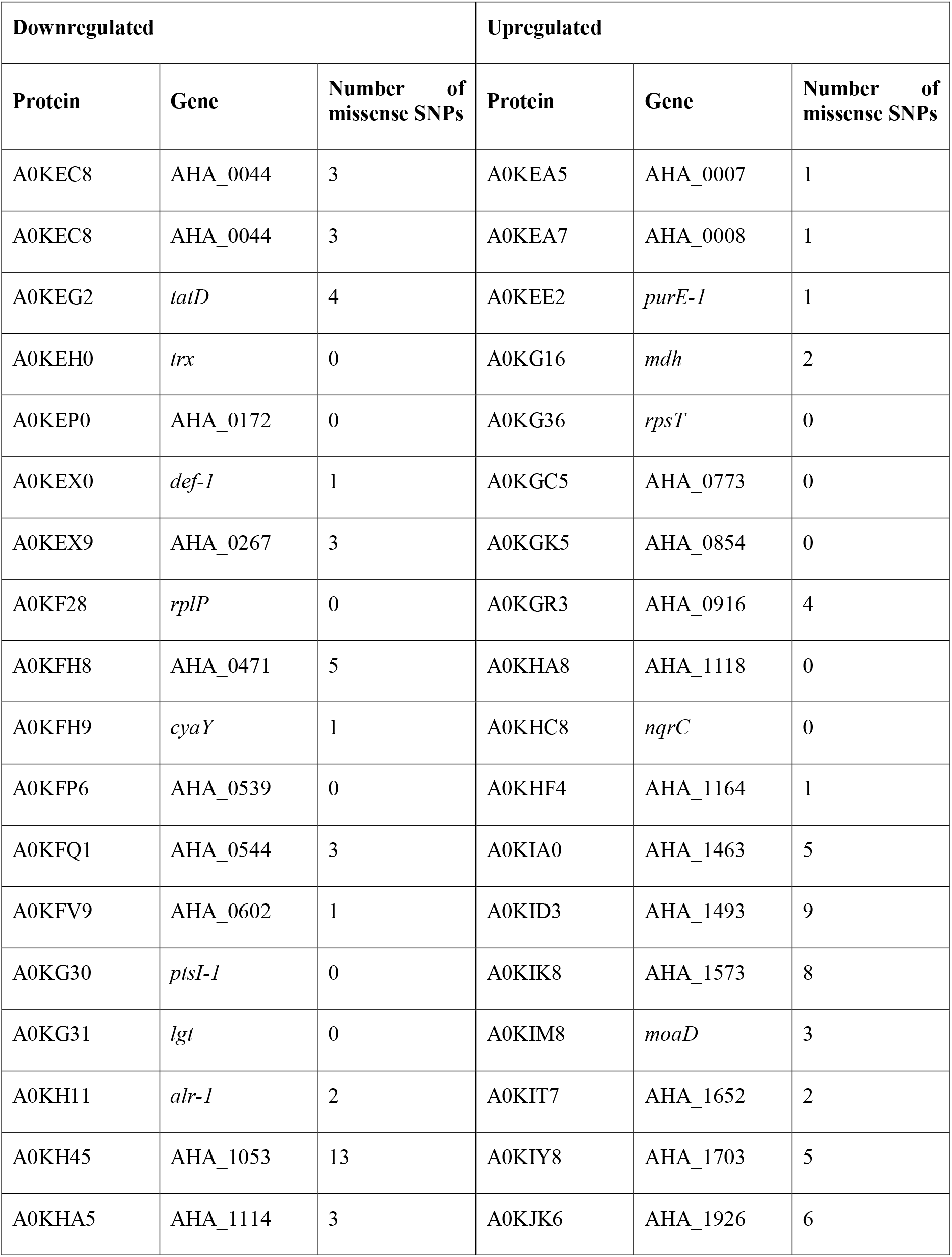
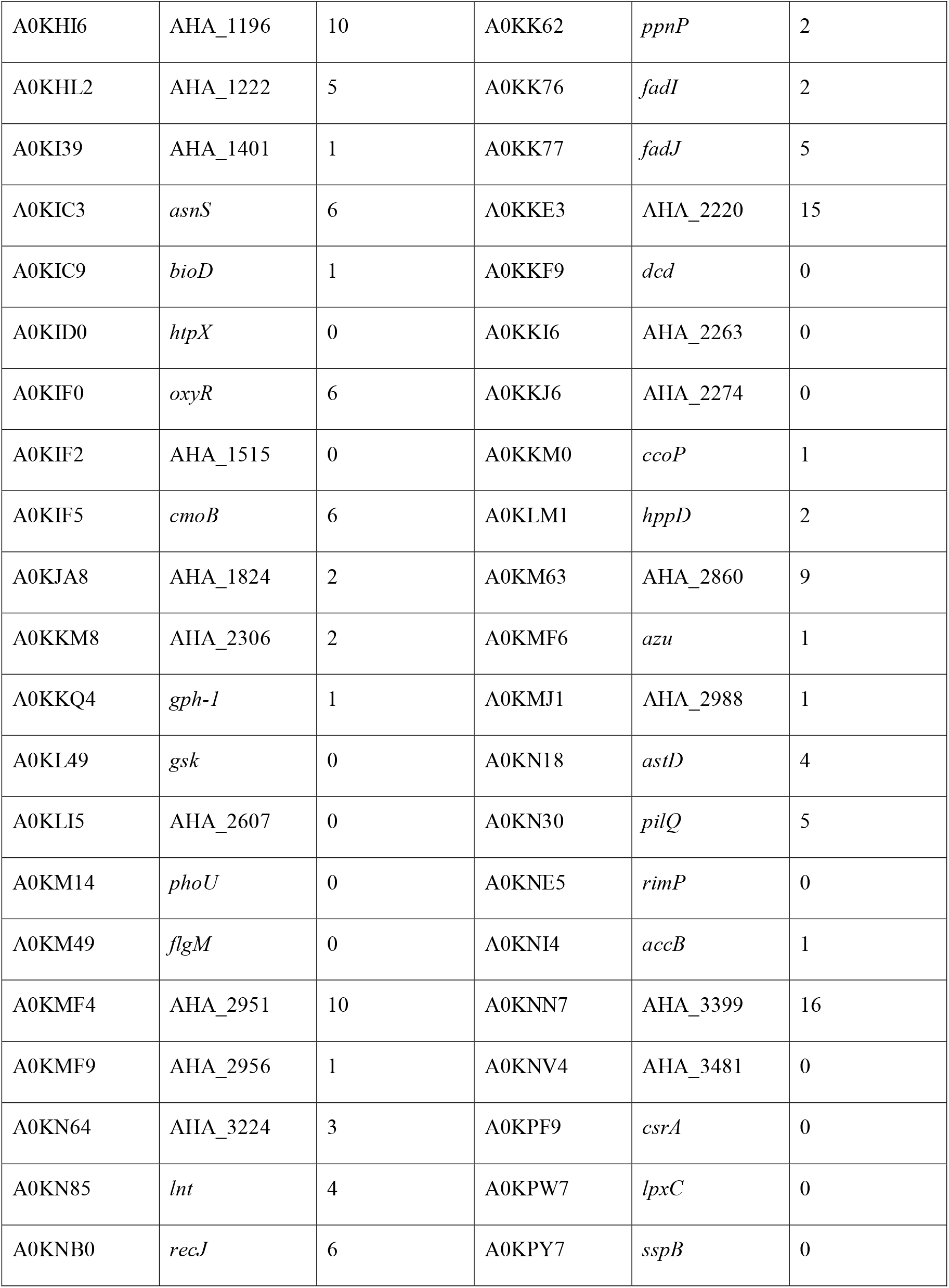
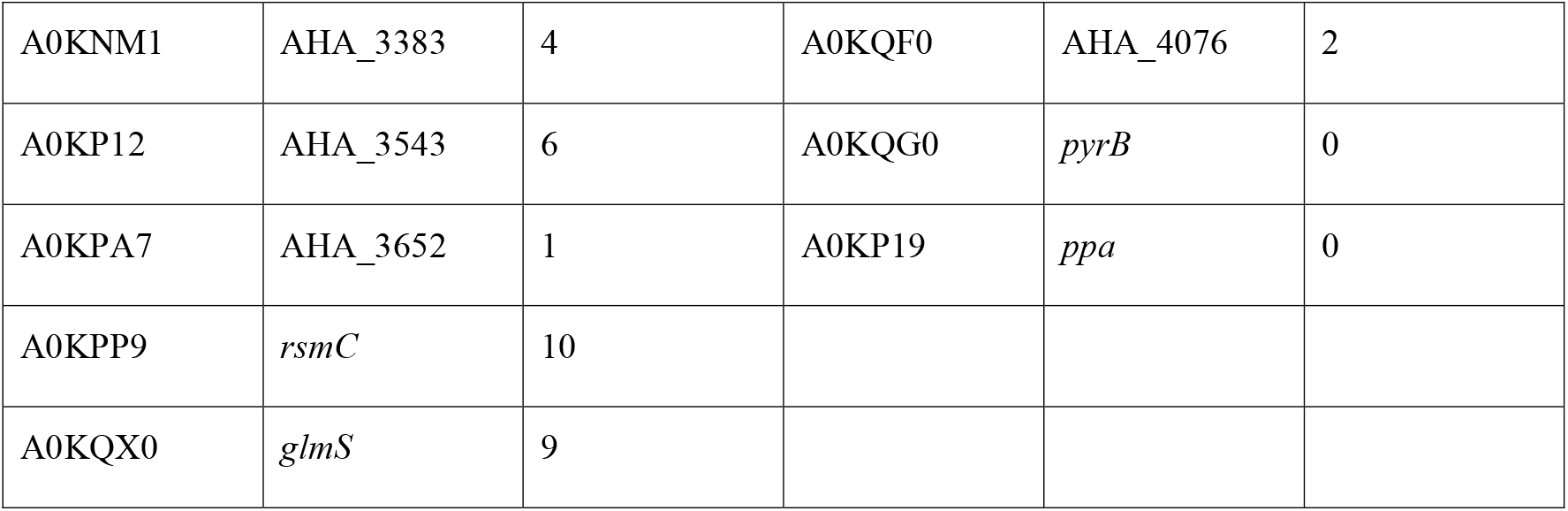
Differential expressed proteins and missense SNPs present in *Aeromonas hydrophila* INISA09.

It is important to mention that since this study was conducted with a pure culture of an isolate bacterial strain under laboratory conditions, it allows us to go deeper into the specific response of one model microorganism, such as *A. hydrophila*, in environments with high BAC concentrations as the activated sludge of a tropical wastewater treatment plant. Nevertheless, we suggest carrying out future studies to evaluate the BAC interactions with other molecules that may be present in the environment, such as antibiotics, and to assess the response in situ conditions or biofilm-forming process to obtain a better panorama of the effect of biocides in microbial organisms from aquatic ecosystems.

## 4 Conclusions

In summary, we characterized, using various genomic, proteomic, and phenotypic techniques, the primary response used by *A. hydrophila* INISA09, an environmental isolate from a Costa Rica domestic activated sludge, to tolerate the presence of a conventional disinfectant as BAC. The strain *A. hydrophila* INISA09 underwent a profound genome rearrangement compared to other strains, probably fine-tuned by thousands of mutations for better survival in the heavily contaminated environment. The low susceptibility of *A. hydrophila* INISA09 to BAC can be related to changes in genes and proteins expression from the outer membrane, transmembrane transport, and fatty acid synthesis metabolic pathways (Figure 6). We provide information on mechanisms used by a environmental multidrug-resistant microorganism exposed to increasing amounts of substances such as disinfectants and other antimicrobials discharged into aquatic systems.

**Figure 6.**
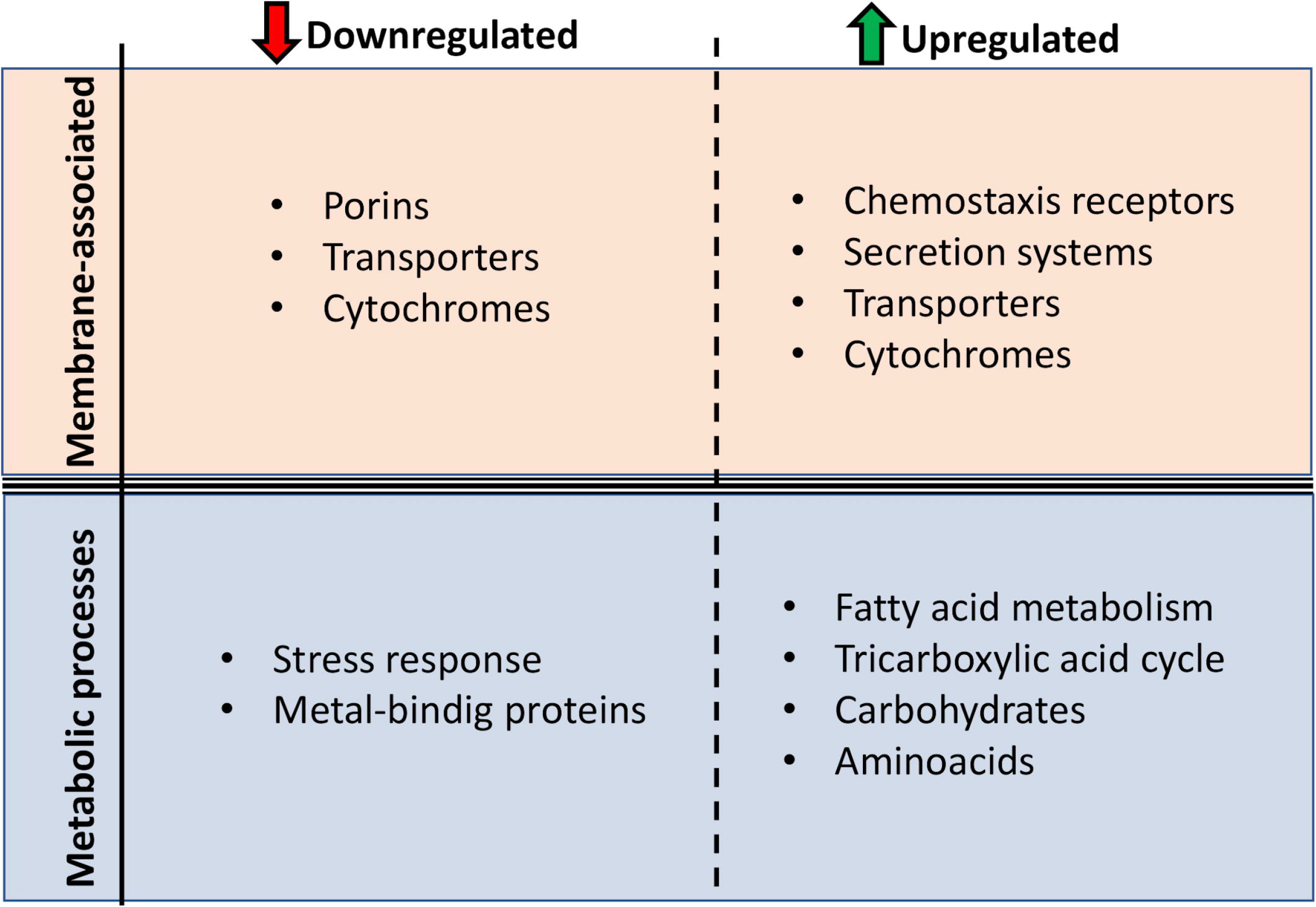
Main protein function changes presented by *Aeromonas hydrophila* INISA09 upon exposure to BAC exposure. The changes were divided in two levels: membrane composition and metabolic processes.

## 6 Author Contributions

LCh, FS, KRJ, and ARR contributed to the conception and design of the study. LCh organized the database. LCh, EGT, and BK performed the data analysis. BK performed the mass spectrometric measurements. LCh, KRJ, and ARR wrote the first draft of the manuscript. LCh, EGT, and BK wrote sections of the manuscript. All authors contributed to the manuscript revision, read, and approved the submitted version.

## 8 Acknowledgments

We thank Dr. Jens Rolff. For mass spectrometry (BK), we would like to acknowledge the assistance of the Core Facility BioSupraMol supported by the Deutsche Forschungsgemeinschaft (DFG).

## 10 Data Availability Statement

The datasets generated and analyzed for this study can be found in the NCBI https://dataview.ncbi.nlm.nih.gov/object/PRJNA896580. All available data are available withing the manuscript and supplementary materials or in repositories when indicted.

